# RaPiD-chamber: Easy to self-assemble live-imaging chamber with adjustable LEDs allows to track small differences in dynamic plant movement adaptation on tissue level

**DOI:** 10.1101/2022.08.13.503848

**Authors:** Ivan Kashkan, Judith García-González, Jozef Lacek, Karel Müller, Kamil Růžička, Katarzyna Retzer, Wolfram Weckwerth

**Author notes:** Equal contribution (Ivan Kashkan, Judith García-González).

## Abstract

Plants rely on fine-tuning organ movement to ensure their survival and productivity. Even subtle loss of directional growth orchestration can result in a huge impact when the plant is impaired to adapt to an ever-changing environment, where it is exposed to manifold exogenous stimuli simultaneously. We present a newly designed chamber to obtain live images to track organ growth and movement differences, called RaspberyPi Dark Chambers (RaPiD-chamber). The RaPiD-chamber is easy to self-assemble and cost-efficient and allows to monitor the continuous growth of etiolated seedlings, as well as their response to light of different wavelengths and from chosen positions. We tested the advice by comparing hypocotyl elongation rate and response to unilateral white and blue light exposure of *Arabidopsis thaliana* Col0. Additionally, we compared the elongation rate of etiolated hypocotyls between Col0 and *kin10*, a mutant lacking the catalytic subunit of the cellular signaling hub SUCROSE NON-FERMENTING RELATED KINASE 1 (SnRK1). *kin10* is known for its diminished ability to control hypocotyl elongation. As a case study, we compared the growth dynamics of etiolated Col0 versus *kin10*. Without further energy source supplementation to the growth medium, the mutant cannot keep up with hypocotyl elongation. Additionally, continuous observation of the dark-grown seedlings allowed us to determine a shift in the dynamics of apical hook angle formation for the mutant.

## 1. Introduction

Plants need to cope with a continuously changing environment ^1–4^. From the very start of plant ontogenesis, even during seed development and germination, plants receive signals from their environment, including information about their position along the gravitropic vector, temperature and light conditions ^5–9^. If the seedling is covered by soil after germination, it initiates hypocotyl elongation until the cotyledons reach light ^10–12^. Once exposed to light, the seedling adjusts the position of cotyledons to optimize photosynthetic light capture and transforms the energy as carbohydrates ^13^. Abundance, localization and metabolism of carbohydrates build the foundation of plant life, survival and productivity, as they function as energy source, signaling molecules and building bricks of cells ^4,14,15^. To understand how hypocotyl elongation, and subsequently its rotation towards light, is modulated on cellular level is of agronomic importance, but how the details of how the molecular network is composed that underpins efficient adaptation is still elusive.

The hypocotyl is a well-established model to study cellular expansion and how morphological changes on the cellular level contribute to organ shape regulation ^16,17^. Cell elongation responses in an everchanging environment occur rapidly, and therefore, to track those dynamic growth adaptations we assembled a low-cost and easy to build up phenotyping device, which we called RaspberyPi Dark Chamber (RaPiD-chamber). This chamber allows to obtain pictures of seedlings germinating and growing on standard growth plates under etiolated or illuminated growth conditions. This allows fast screening of mutant lines or response to additional exogenous triggers such as gravistimulation or pharmacological treatments, to dissect the influence of light responses on plant development and growth. We quantified first growth rate and shape of hypocotyl of etiolated wild type seedlings for several days and afterwards their efficiency to respond to unilateral white or blue light illumination, to test the robustness and quality of our device. Finally, as a case study, we compared hypocotyl growth to test the temporal resolution of the RaPiD-chamber of etiolated Col0 and *kin10*, a mutant with diminished ability to regulate hypocotyl growth rate, carbohydrate metabolism and stress responses under stress conditions.

## 2. Results

### 2.1. Assembling the RaspberyPi Dark Chamber

We present a chamber, called RaspberyPi Dark Chamber (RaPiD-chamber), which allows to obtain live images from seedlings grown on standard square plates used for *Arabidopsis thaliana* seedling cultivation. The pictures can be easily compiled to movies and used for picture analysis to analyze subtle changes in dynamic growth adaptation between different treatments od genetic backgrounds.

A graphical overview of the RaPiD-chamber is shown in Figure 1. Seedlings can be monitored in complete darkness, due to the usage of high-resolution infra-red (IR)-sensitive camera and 880nm IR light-emitting diode (LED) back-light illumination. Plants are considered insensitive towards the light of such wave length, allowing to non-invasively capture organ development and movement of plants grown in total darkness, also known as etiolated seedlings ^17^. Thanks to the red, green, blue and white (RGBW) LED stripes that are positioned left and above the plates harboring the seedlings, plant growth can be monitored under the influence of visible light with different wavelengths (Fig. 2). Also, phototropic responses can be tracked, by illuminating the seedlings only from one side, or together with gravitropic responses when the plate is toppled by 90°. Camera and light sources are connected to the single-board Raspberry Pi 3B+ computer. Imaging process and illumination are flexibly controlled by a Python-based software, which can be configured both remotely and by the intuitive graphical interface via the integrated touch screen (Fig. 1 and 2). Prior to the first imaging cycle, after the plate is placed in a 3D printed holder, the system shots one 16 bit RGB image with the illumination parameters selected by the user, which allows to control the chosen setup remotely without opening the chamber, the material we chose to cover the whole setup. For further imaging, the device uses exclusively the IR LED light source and generates monochrome images in case of the etiolated growth setup, or individually chosen LEDs to study light responses. Pictures are taken within the defined interval and are sent to the remote server to guarantee maximum safety of the experimental data. Obtained images can be analyzed individually or assembled into a stack by the ImageJ software and then used to analyze plant development over time.

**Figure 1:**
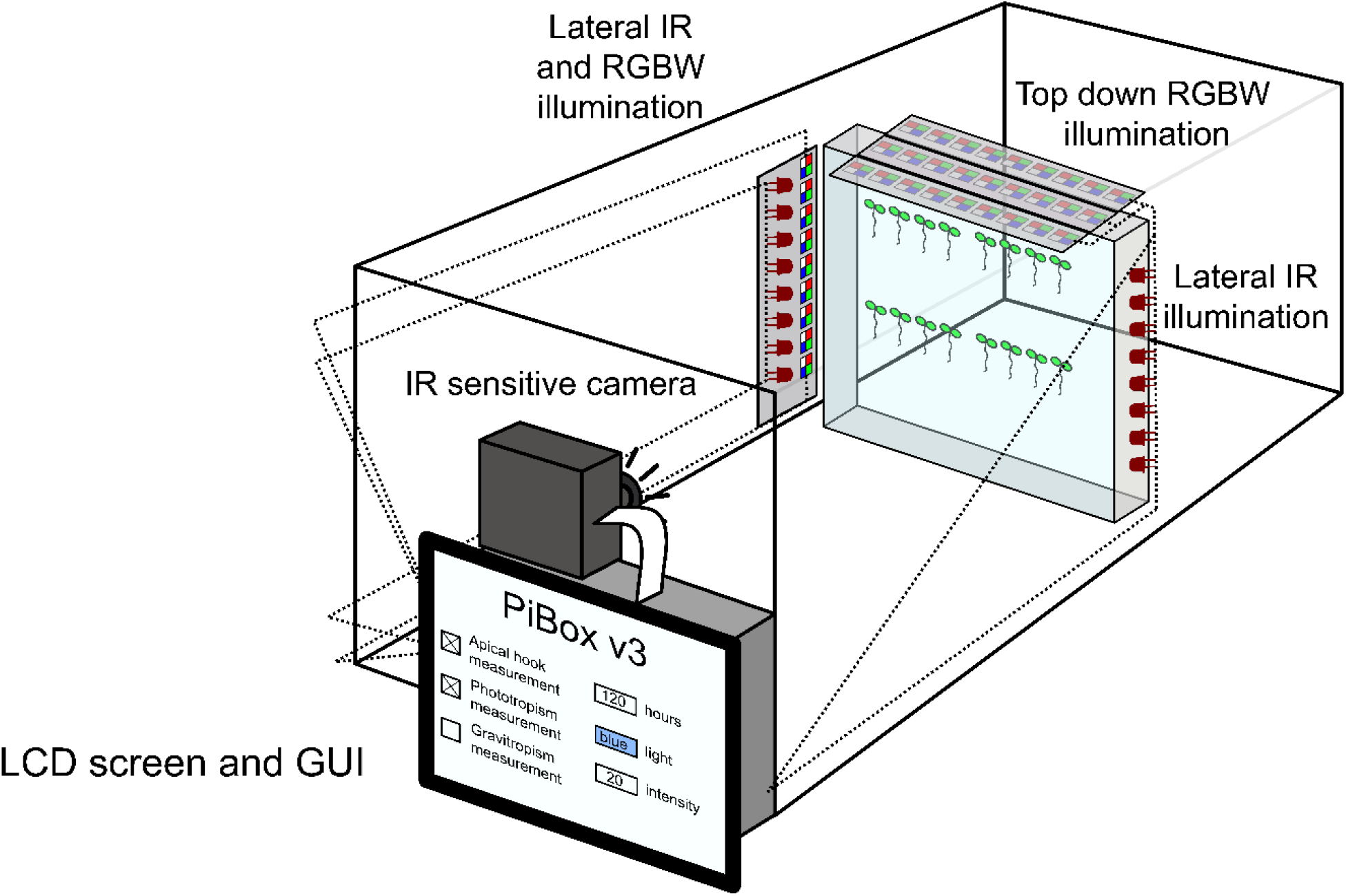
Overview of the RaspberyPi Dark Chamber (RaPiD-chamber). Inside the chamber IR and RGBW LEDs allow to take pictures under different illumination regimes, whereby the direction of the light source is adjustable. Camera and light sources are connected to the single-board Raspberry Pi 3B+ computer, and the pictures are immediately sent wireless to a server, and the experiment can by tracked in real time.

**Figure 2:**
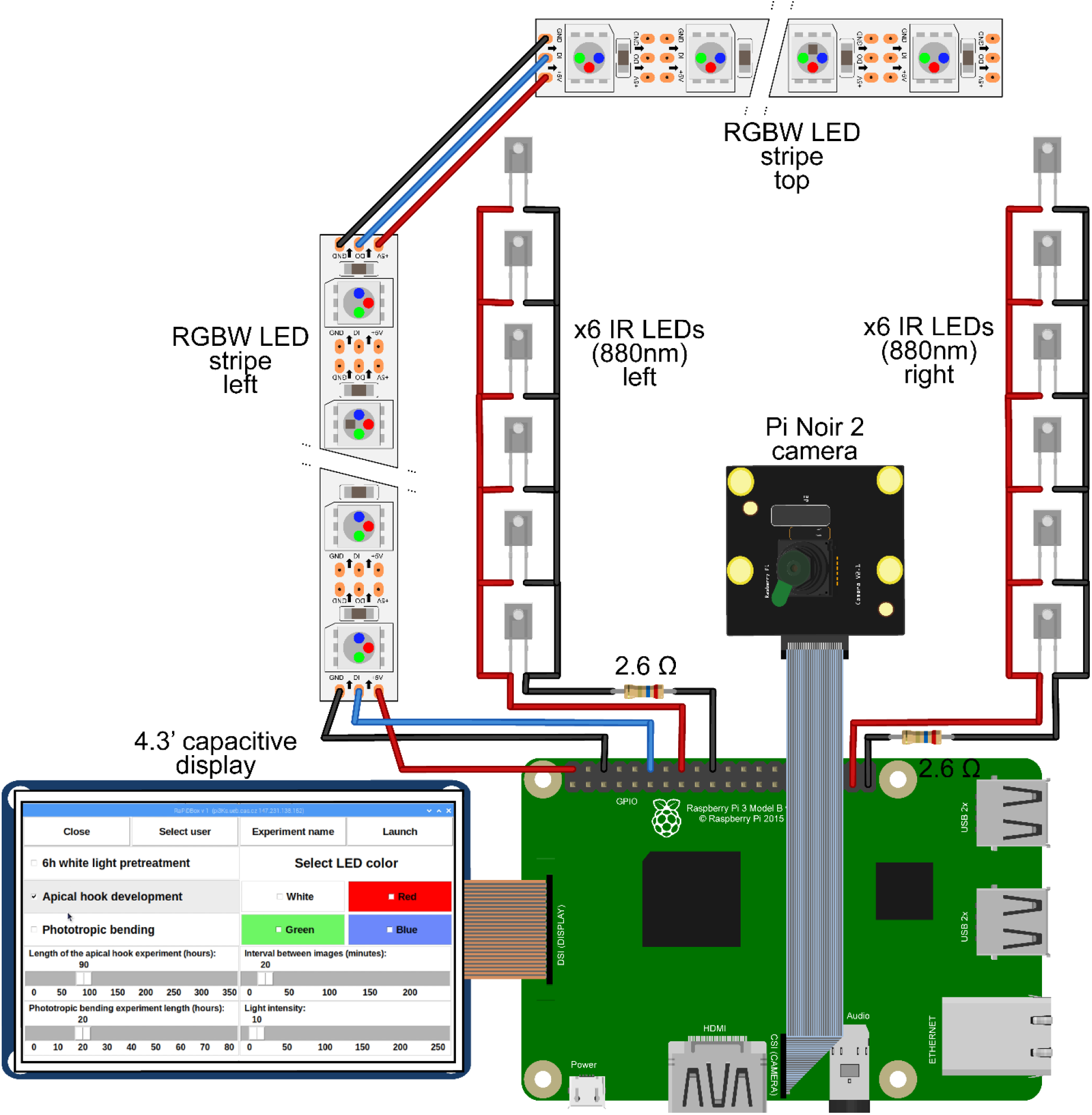
Detailed scheme of camera and led light assembly and wiring. List of used components is summarized in the Methods section.

### 2.2. Robustness of the experimental design

First, we tested the robustness of our experimental setup to be sure that we obtain the same rage of responses over long time periods and compared the growth curves to previously published studies ^17–20^. To this end, we assembled individual RaPiD chambers to run experiments in parallel, were placed in a thermoregulated growth chamber at 21 °C. To evaluate hypocotyl growth traits of seedlings germinated in complete darkness, we plated Col0 seeds on ½ MS medium, without additional sugar supplementation, stratified the seeds for 48h at 4 °C and exposed them for 6h to light before placing the plates in the RaPiD-cambers. Hypocotyl elongation and apical hook growth occurred as previously described ^17^. We tracked growth traits including hypocotyl length in mm (Fig. 3A), growth rate in mm.h^-1^ (Fig. 3B) and apical hook angle in degrees (Fig. 3C), over a scope of 65 Hours After Germination (HPG), from the moment the hypocotyl was visible around seven HPG.

**Figure 3:**
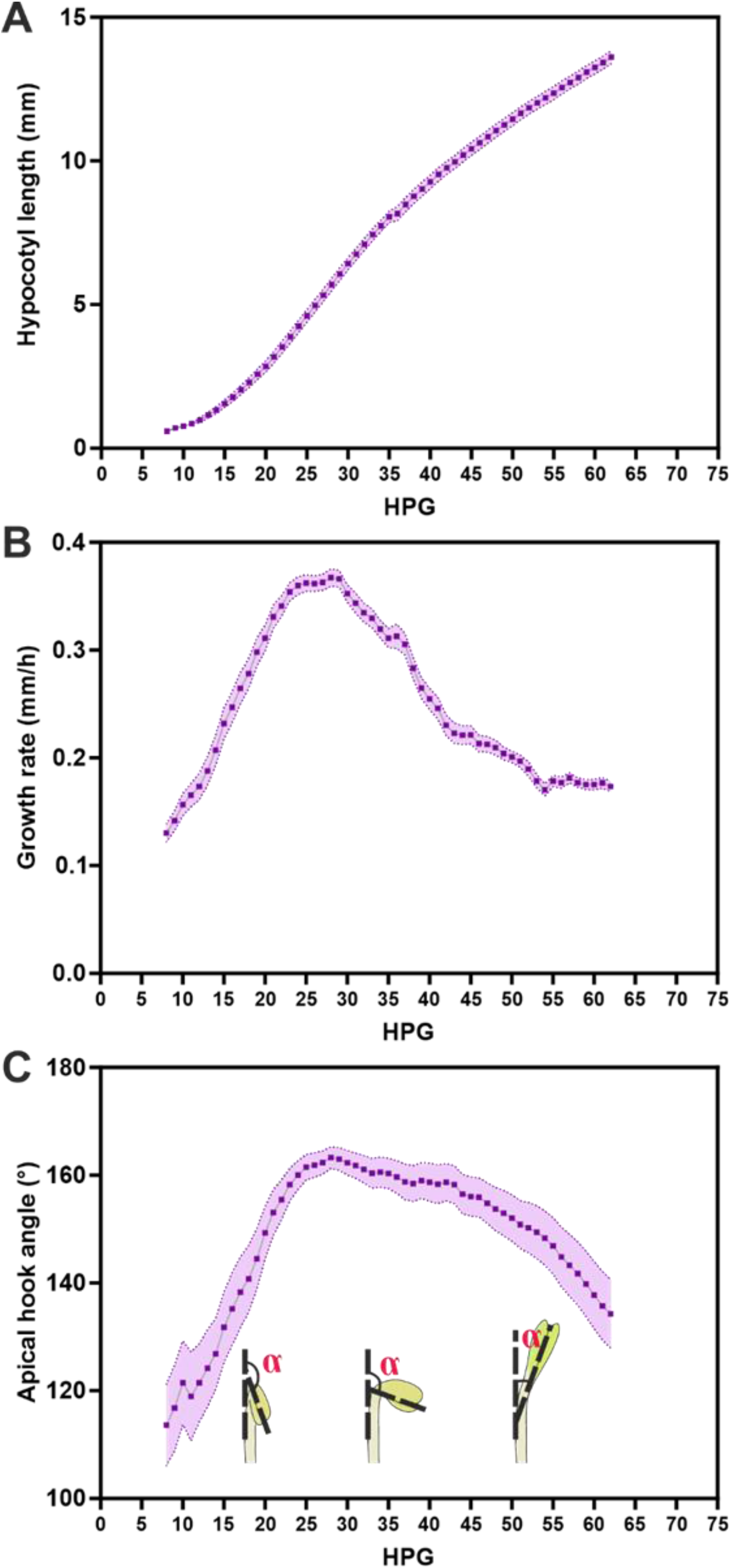
Etiolated hypocotyl growth dynamics of wildtype (Col0) seedlings from 7 to 62 Hours after Germination (HPG). Growth kinetics were defined by: A, Hypocotyl total length. B, Hypocotyl elongation rate in mm.h-1. As expected, with progressing duration of growth in total darkness elongation rate declines stepwise. C, Apical hook angle as described in the inset diagram. After a brief formation phase till around 25h, the hook is maintained to protect the meristematic tissue, and opens gradually after around 45h in our setup. The timing of individual steps of apical hook development resembles results previously published, for example by Zhu et al., 2016 ^17^ and appeared constant among individual biological repetitions, wherefore we consider our experimental setup as robust. Each data point represents the mean of 59 seedlings measured in a total of 2 biological replicates. Shaded bands represent standard error.

Hypocotyls elongate until they reach maximum length that is limited by the plant internal energy supply that was stored in the seed ^21^, as we are not supplementing the growth medium with sugar. Exogenous sugar supplementation is enhancing elongation rate of hypocotyls and may result in masking phenotypes ^22–25^. During ongoing germination in darkness, the apical part of the hypocotyl forms a hook like structure, which is meant to protect the meristem from mechanical harm while the hypocotyl pushes through the soil ^26^. After the formation and maintaining phases, the apical hook undergoes an opening phase, which will allow cotyledon unfolding and positioning towards light source ^17,26^. All phases of apical hook maturation require spatially and temporally precise regulation of asymmetric cell elongation at opposite position of the hypocotyl, and although several molecular key players were identified that underpin asymmetric growth along the apical-basal axis, the RaPiD-chamber will allow to dissect the temporal aspect of their action further. Although total hypocotyl length increases over time, until all plant stored resources are used up, the elongation speed declines around 45h after HPG, which matches with the curve we obtained measuring the dynamics of apical hook opening ^19^.

### 2.3. Testing the illumination system by tracking Col0 phototropism response to unilateral white versus blue light exposure

The improvement of our device compared to already existing systems ^17^ is the implementation of light sources emitting different wavelengths, which can be placed according to the users’ needs in different positions. To test our illumination setup, we compared the impact of unilateral white and blue light on cotyledon positioning and dynamics of phototropic response towards the light source (Fig. 2). Phototropic responses that result in the reorientation of the cotyledons towards the light source were described to be much more sensitive and mechanistically distinct from general light induced hypocotyl growth inhibition and apical hook opening responses compared to growth conditions when etiolated seedlings are evenly exposed to light ^19^. Whereby on one side, direct illumination of the hypocotyl stops it elongation and enhances apical hook opening, unilateral illumination promotes asymmetric elongation at opposing sides to initiate bending of the hypocotyl towards the light source ^19^. We compared the impact of white versus blue light, because monochromatic blue light on its own stimulates positive hypocotyl phototropism, but with slight shifted kinetics compared to white light, where red light and the ratio of red to far-red light influences different aspects of hypocotyl growth ^27–29^.

As previously published ^19^, illumination with monochromatic blue light has stronger reducing effect over time on hypocotyl elongation compared to white light illumination (Fig. 4A). Phototropic response of the hypocotyl results first in elevatedelongation rate to reach the light source, and then declines when the cotyledons are properly positioned (Fig. 4B). The modulation of elongation speed upon unilateral illumination is reflected in the extent of curvature of the bending hypocotyl towards the light source (Fig. 4C). During the period of elevated elongation, a curvature deviates, and when the cotyledons are reoriented, the elongation rate declines and also bending is terminated (Fig. 4C, D). Monochromatic blue light has a stronger impact on hypocotyls phototropic response, which is also visible in the enhanced curvature upon blue light compared white light exposure (Fig. 4C). Overall, monochromatic blue light perception leads to both, phototropic curvature enhancement and inhibition of stem elongation in higher plants ^30,31^. This would explain, why after a more pronounced curvature of the hypocotyl, and afterwards a rapid reduction of the elongation rate of seedlings responding to unilateral blue light (Fig. 4B, C). Illumination of the etiolated seedling results furthermore in enhanced apical hook opening until the angle reaches almost 0°, when the seedling is evenly illuminated, to ensure the best possible orientation of the cotyledons towards the light source to maximize phototropic activity ^18,19^. Overall, our illumination setup is suitable to track growth adaptations upon changing light regimes.

**Figure 4:**
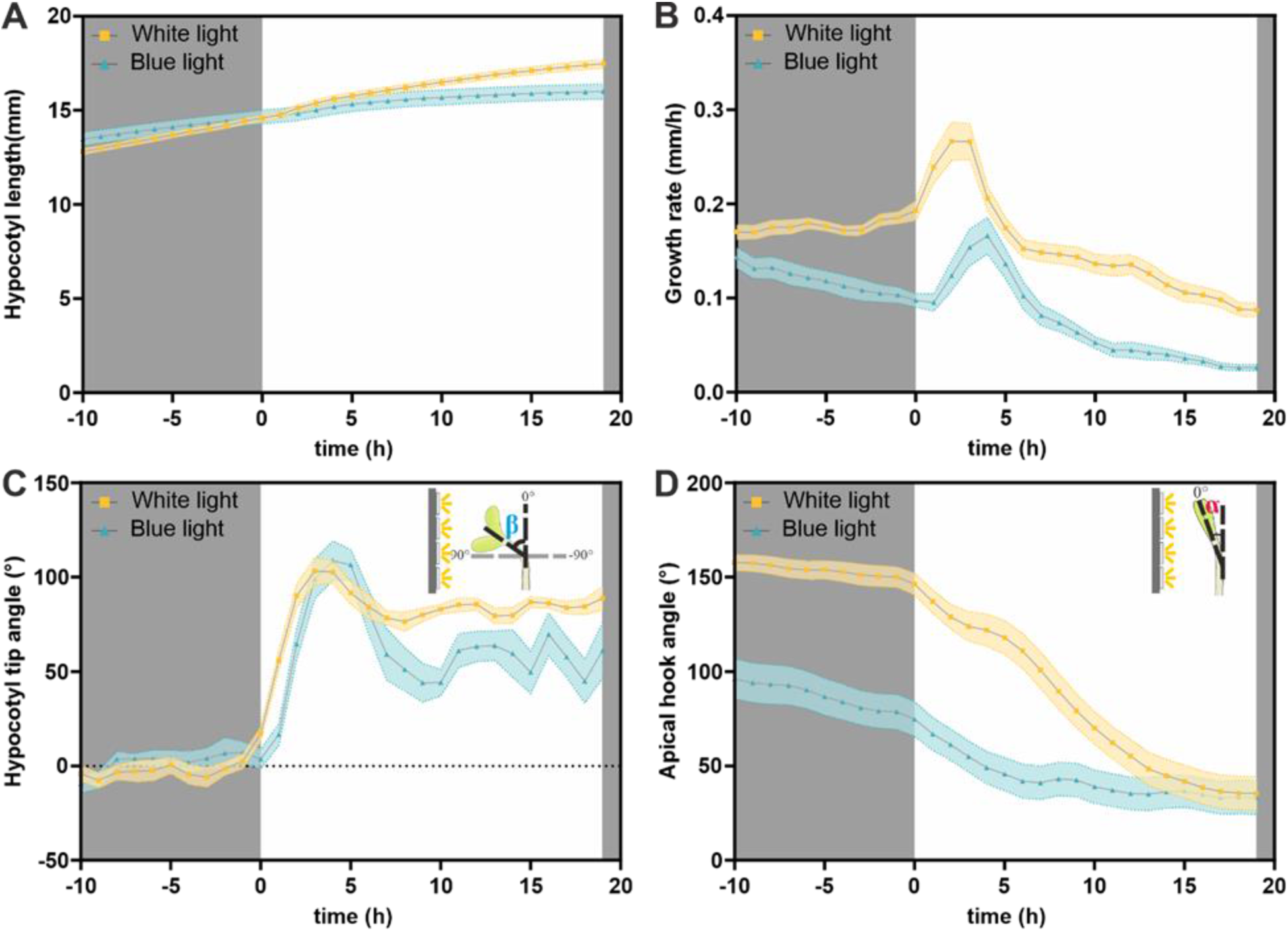
Comparison of phototropic response of etiolated seedlings towards white light versus monochromatic blue light. A, Quantification of hypocotyl length over time, starting with etiolated seedlings at the age of 52 h HPG, till 19h after unilateral illumination from the left side. B, Quantification of the overall growth rate in mmh-1 of the same seedlings. C, Hypocotyl tip angle reflects curvature formation during phototropic response towards the light source. D, Evaluation of the apical hook angle during the positive phototropic response of the hypocotyl towards the unilateral light source. Each data point represents the mean for 19 to 24 plants, shaded bands represent standard error.

### 2.4. Continuous tracking of a mutant impaired in hypocotyl growth rate regulation and carbohydrate metabolism revealed so-far overlooked phenotypes of dark germinating seedlings

The seed stores energy in form of starch, which is utilized during germination and early seedling development ^21^. If the seedling germinates being shaded from light, the energy resources are used to power hypocotyl elongation, while other developmental and growth processes are reduced to ensure the cotyledons will reach light in time ^32,33^. Once exposed to light, photosynthesis in the cotyledon generates enough energy to change developmental program from skotomorphogenesis to photomorphogenesis ^13,34^. Every adaptation to environmental conditions results in reorganization of plant architecture, underpinned with energy costly adjustment at proteomic and metabolomic level ^4,14,35–37^. Plants integrate energy and nutrient status to regulate growth and stress responses using antagonistic signaling pathways ^3,38–43^. The molecular master regulator complex SUCROSE NON-FERMENTING RELATED KINASE 1 (SnRK1) plays a fundamental role orchestrating anabolic and catabolic signaling cascades that underpin plant morphological changes upon starvation, stress signaling, but also for normal growth and development where resource availability needs to be balanced with the sum of external and internal stimuli ^14,37,43–45^. The SnRK1 complex is an evolutionary highly conserved energy sensor, composed of three subunits, the catalytic-α and regulatory β- and γ-subunits ^44,46–48^. The Arabidopsis thaliana genome encodes two functional catalytic α-subunit isoforms, SnRK1α1 (KIN10; At3g01090) and SnRK1α2 (KIN11; At3g29160), and whereas both are expressed in many tissues and developmental stages redundantly, SnRK1α1 is responsible for most of the kinetic activity of the SnRK1 complex ^44,47,49,50^. Therefore, we used in our case study a knockout mutant of SnRK1α1/KIN10 to compare its performance during germination in complete darkness regarding hypocotyl length, hypocotyl elongation rate and apical hook development.

With reduced access to light the plant enhances hypocotyl elongation to enhance the probability to reach more illuminated regions, and in previous hypocotyl elongation assays *kin10* seedlings had shorter hypocotyls compared to Col0 when grown on medium without sugar supplementation ^22,23^. Continuous monitoring of hypocotyl elongation rate confirmed the strong diminished hypocotyl length (Fig. 5A) and growth rate (Fig. 5B) of *kin10* compared to Col0. Furthermore, the immediate monitoring of the seedlings from germination start onwards, shows that growth deficiency is accumulating over time, till it peaks during the timeframe of apical hook maintenance phase, and afterwards declines steep for the wild type seedling, and aligns with the curve of kin10 elongation rate, which correlates with the pattern of apical hook opening phase (Fig. 5A, B). When we correlate the apical hook angle with the growth rate, we see that during the apical hook maintenance phase, where we quantified the biggest difference between Col0 and *kin10* for both traits (Fig. 5B -D).

**Figure 5:**
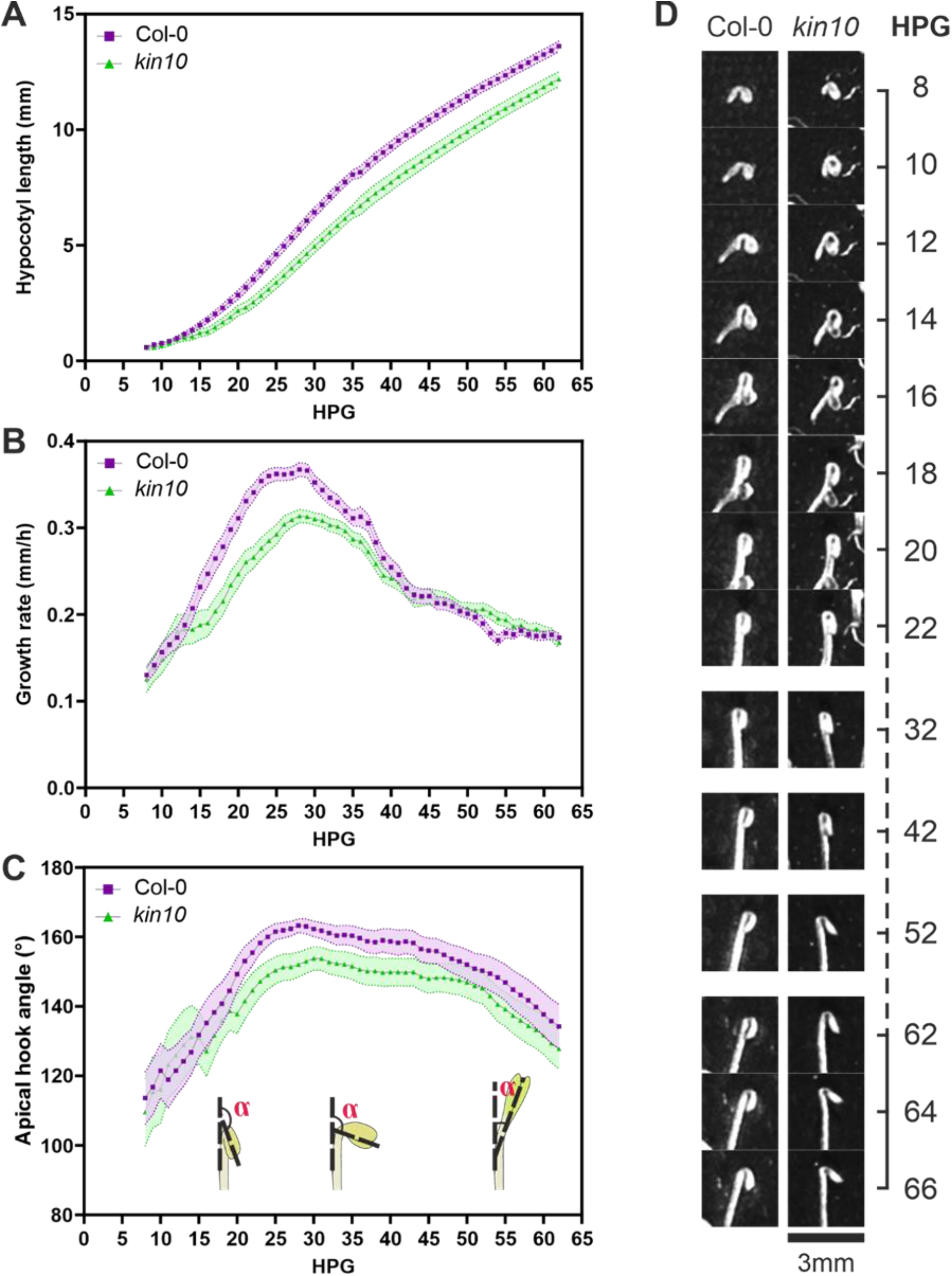
Comparison of hypocotyl growth traits of etiolated Col0 versus kin10 seedlings. Seedlings were tracked from 7 to 62 HPG. A, Quantification of hypocotyl length in mm. B, Quantification of growth rate in mmh-1 C, Apical hook angle formation. Each data point represents the mean of 54 to 59 seedlings. Shaded bands represent standard error. D, Representative images depicting multiple timepoints in hook development of the studied lines.

Because SnRK1 orchestrates manifold cellular processes, and several studies described its regulatory role in starch deposition as well in degradation, we performed Lugol’
ss staining of the etiolated seedling at the end of the experiment and compared cell appearance and starch localization in the hypocotyls of Col0 and kin10. The etiolated kin10 seedlings show abnormal cell morphology, and that the cells cannot fully expand, whereas Col0 hypocotyl cells are apparently longer (Fig. 6). Furthermore, the mutant displays a high accumulation of stained starch granule (Fig. 6B). Further studies are required to dissect if this diminished starch turnover results in changed cellular mechanoproperties of hypocotyl cells, or to which extend carbohydrate signaling and energy homoeostasis are interconnected with other signaling pathways to regulate growth rate and apical hook formation is required.

**Figure 6:**
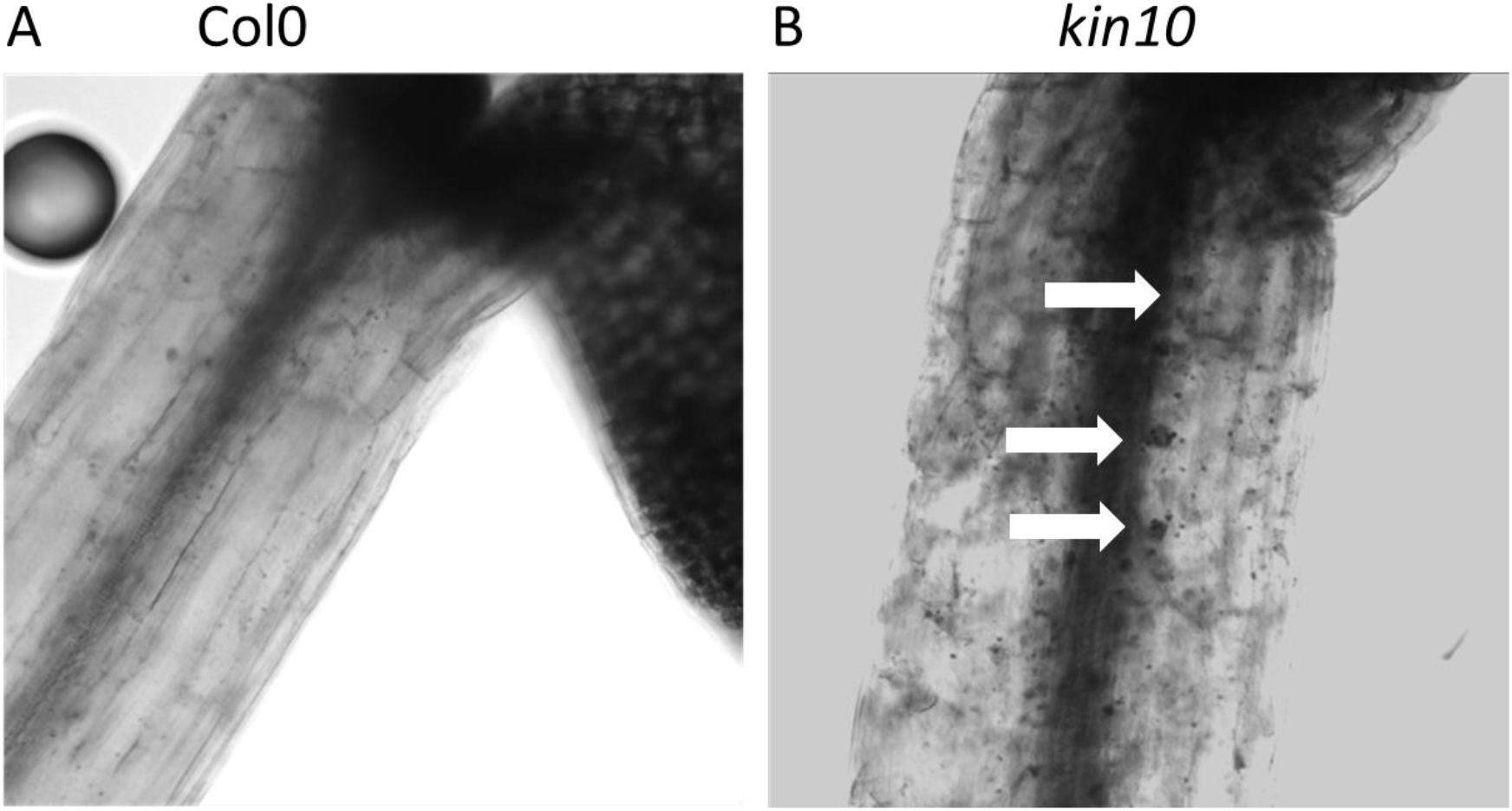
Lugol’
ss staining of etiolated Col0 and *kin10* seedlings. Brightfield images of the apical part of the hypocotyls, focus on epidermal cells, of 7 DAG old etiolated seedlings. Hypocotyls of kin10 show dramatic cell morphological changes and enhanced accumulation of starch granule (indicated by white arrows).

## 3. Methods

### 3.1. Material used to assemble the RaPiD chamber

Single-board computer Raspberry Pi 3B+ (Raspberry Pi Foundation, Cambridge, UK) was used as a computing module. For the imaging in the dark, Raspberry Pi Camera Module 2 with removed IR filter (Raspberry Pi Foundation, Cambridge, UK) was used. For the back-light illumination we used two stripes of OSIXCA5111A 880nm IR LED’s (OptoSuply limited, Hong Kong, CN) connected in parallel. For the uni-lateral and top-down illumination we used one 240 mm stripe of 28 SK68121M144RGBNWW30 LED’s (BTF-LIGHTING Teachnology Co., Limited, Guandong, CN). Waveshare 4,3” DSI LCD capacitive touch display (Waveshare electronics, Guandong, CN) was used for the visualization of the graphic user interface (GUI). The exact wiring scheme, Raspberry Pi setup manual and code used in the RaPiD assembly can be found in the GitHub depository: https://github.com/lamewarden/RaPiD-boxes-software.

### 3.2. Plant material and growth conditions

Seed stock of Col-0 was obtained from the Laboratory of Hormonal Regulations in Plants, Institute of Experimental Botany, Czech Academy of Sciences and *kin10* from NASC collection and tested for insertion prior experiments. Verification of mutation in At3G01090 was done by PCR amplification using GoTaq polymerase (Promega) at annealing temperatures 56°C and 62°C and primers specific to Kin10 (At3G01090_forward: 5’- AGCAAGAAAAGCAGCACACA-3’, reverse: 5’- GTCGCCCAATGTTGTCAAGT-3’). As a control, primers specific to Ims2 were used (At5G23020_foward: 5’- GGCATTCACCAGGTTCCATA-3’, reverse: 5’- TCTATTGTGCCATGGTCCCA-3’). 20x diluted genomic DNA isolated from seedlings of Col-0 and GABI579E09 plants using DNeasy Plant Mini kit (Qiagen) was used as a template.

Seeds were surface sterilized using 50% (v/v) bleach and 0.1% Tween20 (Sigma-Aldrich, Darmstadt, Germany) for 5 min and then rinsed three times with sterile water. The seeds were plated on ½ Murashige and Skoog (Sigma) medium, solidified with 1% agar (Sigma) and adjusted to pH 6.0 by KOH. The seeds were plated and stratified at 4 °C for two days and germination was induced by placing the seeds in 100 ^μmol m−2 s−1^ white light for 6 h. Afterwards the plates were placed in the RaPiD-chamber and monitoring was immediately started. Seedlings germinated approximately 28h after their transfer to the chamber, and 7 HPG (Hours Post-Germination) were required to track first differences in the elongation rate of etiolated hypocotyls and apical hook opening between Col0 and kin10, and secondly for the phototropism experiment a unilateral white light source was turned on at 62HPG and we recorded the response for further 19 h.

### 3.3. Data evaluation and plot preparation

Hypocotyl traits were analyzed using the freely available Fiji software ^51^. Hypocotyls were evaluated individually using the multipoint tool to obtain their tip coordinates at each timepoint of the image series. Apical hook was measured using Fiji’s built-in angle tool. Basic data manipulation was done using MS Office Excel.

### 3.4. Lugol’ ss staining

Seven-day old etiolated seedlings, grown on ½ MD medium without sugar supplementation, were stained with Lugol iodine solution for microscopy (Sigma, 62650-1L-F) according to the manufacturer’s description. We captured images with brightfield illumination at the Zeiss 880 microscope with EC Plan-Neofuar 20x/0.50 (WD=2.0 mm) objective, with focus on the epidermis of the apical part of the hypocotyls.

## 4. Discussion

We present a low-cost and easy to self-assembly monitoring device of seedlings grown under different illumination regimes, including the position of the light source. The advice can be assembled in a card box, so the height and distance to camera can be easily adjusted to the research requirements, and can stay small enough, so it can be placed on any shelf in a growth chamber. The LEDs, camera and controls are available in electronic stores and the setup is fast assembled. Direct, wireless transmission of the pictures, whereby intervals and duration of experiments can be changed remotely, to a server allow to control or adjust the experiment from any place with internet connection. The RaPiD-chambers can be used to efficiently phenotype even subtle changes in organ growth dynamics, depending on light quality. We confirmed the robustness of the device, and compared the growth dynamics of *Arabidopsis thaliana* wild type Col0 and a mutant impaired in hypocotyl expansion to obtain differences in their growth dynamics in not yet available temporal resolution.

Hypocotyls are an optimal study object to decipher so-far elusive connection points of signaling cascades and role of molecular key players that orchestrate tropistic responses in dependence on energy and nutrient level of the plant, which to a big extent are modulated over SnRK1 activity ^4,44,46,47,52^. Because, hypocotyls consist of constant number of cells with high elongation ability, the influence of exogenous stimuli can be easier correlated to cell expansion responses depending on nutrition conditions and genetic background ^10,25^. Epidermal elastic asymmetry is required to ensure efficient and fast directional hypocotyl growth and drives anisotropic growth of the rapidly elongating cells ^33^. It is of high interest how establishment of cellular mechanical traits and well-characterized phytohormonal signaling pathways are modulated during hypocotyl elongation and apical hook establishment by SnRK1. A recent study showed the role of SnRK1 in regulating key catabolic genes required for stored resource mobilization during early seedling development ^21^. As a central cellular hub for signaling cascades upstream of cell morphology modulation, SnRK1 could for example orchestrate the interplay of molecular networks connecting carbohydrate metabolism and hormonal signaling, which includes the auxin-callose feedback circuit, which is required for efficient hypocotyl growth regulation ^53^.

Altogether, the application of the RaPiD-chamber allowed us to phenotype hypocotyl growth responses at high temporal resolution. Beside the demonstrated growth conditions in this study, it is further possible to observe seedlings grown under full illumination, with white light or under individual wavelengths. Additionally, gravitropic response, after changing the orientation of the cultivation plate by 90°, can be monitored. The low costs and easy assembling of the chamber allow to establish a phenotyping unit to track continuously dynamic plant growth responses, which allows to reveal differences on tissue level that would be overlooked by comparing end-point or occasional scans of seedlings. Finally, because the plates are stationary, no unnecessary movement of the seedlings is interfering with the outcome of the experiments.

## 6. Further information

### Funding

This research was funded by the Ministry of Education, Youth and Sports of Czech Republic from European Regional Development Fund ‘Centre for Experimental Plant Biology’: Projects no. CZ.02.1.01/0.0/0.0/16_019/0000738 for K.Re and K. Rů., 19-23773S for K.Rů., and MEYS CR - LM2018129 Czech-BioImaging for I.K.

### Author Contributions

Conceptualization, I.K., W.W. and K.Re.; methodology, I.K., J.L., K.M., K.Re. and J.L.; validation, K.Re. and J.L.; formal analysis, J.G.-G.; investigation J.L., K.Re. and J.L.; resources, K.Re and K. Rů.; data curation, K.Re. and J.L.; writing—original draft preparation, I.K., K.Re. and J.G.-G.; writing—review and editing, W.W., K.Re., J.G.-G. and K. Rů..; visualization, J.G.-G. and K.Re.; supervision, K.Re.; project administration, K.Re.; funding acquisition, K.Re and K. Rů. All authors have read and agreed to the published version of the manuscript.

